# WDR45 contributes to neurodegeneration through regulation of ER homeostasis and neuronal death

**DOI:** 10.1101/282210

**Authors:** Huida Wan, Qi Wang, Xiuting Chen, Qiufang Zeng, Yanjiao Shao, Houqin Fang, Xun Liao, Husong Li, Minggang Liu, Tian-Le Xu, Dali Li, Minyao Liu, Bo Meng, Bin Tang, Zhuohua Zhang, Lujian Liao

## Abstract

Mutations in the autophagy gene *WDR45* cause β-propeller protein-associated neurodegeneration (BPAN); however the molecular and cellular mechanism of the disease process is largely unknown. Here we generated constitutive *Wdr45* knockout (KO) mice that displayed cognitive impairments, abnormal synaptic transmission and lesions in hippocampus and basal ganglia. Immunohistochemistry analysis shows loss of neurons in prefrontal cortex and basal ganglion in aged mice, and increased apoptosis in these regions, recapitulating a hallmark of neurodegeneration. Quantitative proteomic analysis shows accumulation of endoplasmic reticulum (ER) proteins in KO mouse. Furthermore, we show that a defect in autophagy results in impaired ER turnover and ER stress. The unfolded protein response (UPR) is elevated through IRE1α and possibly other kinase signaling pathways, and eventually leads to neuronal apoptosis. Suppression of ER stress, or activation of autophagy through inhibition of mTOR pathway rescues neuronal death. Thus, our study not only provides mechanistic insights for BPAN, but also suggests that a defect in macroautophagy machinery leads to impairment in selective organelle autophagy.

## Introduction

Macroautophagy (herein referred to as autophagy) is the major cellular catabolic process to degrade damaged organelles as well as protein aggregates. The process involves the engulfment of cellular compartments by a double-membrane structure and the fusion with lysosomes for degradation (Choi, Ryter et al., 2013). The importance of autophagy in preventing neurodegeneration has been greatly appreciated. In mice, deletion of the essential autophagy gene *Atg5* or *Atg7* in the central nervous system results in accumulation of intracellular inclusion bodies and neurodegeneration (Hara, Nakamura et al., 2006, Komatsu, Waguri et al., 2006). In humans, mutations in various genes functioning in multiple steps of autophagy cause a wide range of neurodegenerative diseases (Menzies, Fleming et al., 2017, Menzies, Fleming et al., 2015, Yamamoto & Yue, 2014).

β-Propeller protein-associated neurodegeneration (BPAN), also known as static encephalopathy of childhood with neurodegeneration in adulthood (SENDA), is a recently identified subtype of neurodegeneration with brain iron accumulation (NBIA). BPAN is characterized by global developmental delay in motor and cognitive function during early childhood and remains static. During early adulthood the patients develop sudden-onset dystonia, Parkinsonism, and dementia (Araújo, Garabal et al., 2017, Gregory & Hayflick, 2011, Kasai-Yoshida, Kumada et al., 2013). To date, most studies on BPAN have focused on case reports and genetic analyses, confirming BPAN as a distinct class of neurodegenerative disease. Recently, *de novo* mutations in *WDR45* gene were identified in BPAN patients by exon sequencing (Haack, Hogarth et al., 2012, Saitsu, Nishimura et al., 2013, Wynn & Pulst, 2017). Further studies on patient-derived lymphoblastoid cells show lower autophagic activity and accumulation of early autophagic structures (Saitsu et al., 2013), linking an abnormal autophagy to the disease.

Structured as a β-propeller-shaped scaffold protein, WDR45 (also known as WIPI4) is one of the four mammalian homologs of yeast Atg18, which plays important roles in autophagy (Bakula, Müller et al., 2017). These include recruitment of lipids via its phosphatidylinositol-3-phosphate (PtdIns3P)-binding ability and regulation of autophagosome formation via interacting with ATG2 and AMPK/ULK1 complexes (Bakula et al., 2017, Dall’Armi, Devereaux et al., 2013, Tooze & Yoshimori, 2010). Unlike other essential autophagy genes such as *Atg1, Atg5*, and *Atg7*, whose germ line deletion is lethal; germ line deletion of *epg-6*, the worm homolog of WDR45, in *Caenorhabditis elegans* only causes abnormal autophagic structures (Lu, Yang et al., 2011). Neuronal-specific knockout of *Wdr45* in mouse results in dysfunction in autophagy and axonal degeneration, together with poor motor coordination and impaired learning and memory (Zhao, Sun et al., 2015). Thus, it appears that the major function of WDR45 is the regulation of autophagosome formation and its loss-of-function could contribute to behavioral abnormalities in mice.

In spite of these studies, it remains a mystery how loss of WDR45 leads to neurodegeneration in BPAN, and whether dysfunction in autophagy is the major cellular underpinning of the disease. To address these questions, we generated constitutive *Wdr45* knockout (KO) mouse to model the complex processes of BPAN. Multiple levels of analyses show that the KO mouse displayed cognitive impairments, abnormal synaptic transmission and lesions in hippocampus and basal ganglia. Comparative proteomic quantitation of brain regions shows a surprisingly large number of accumulated endoplasmic reticulum (ER) proteins in KO mouse. Mechanistically, we find that defects in autophagy results in impaired ER turnover and ER stress even under basal conditions. The unfolded protein response (UPR) is augmented under basal conditions and dramatically aggravated in KO cells after inducing ER stress, which leads to induction of neuronal apoptosis. These abnormalities can be rescued by mTOR inhibitor as well as by inhibition of UPR. Thus, our study pinpoints the molecular mechanism on how a dysfunction in general autophagy machinery leads to organelle selective autophagy, which leads to BPAN-like pathology. These data also support a unified principle explaining the onset of neurodegeneration in BPAN patients.

## Results

### *Wdr45* knockout mouse displays cognitive impairments

We generated *Wdr45* KO mouse using CRISPAR-Cas9 technique targeting the mutated sites found in SENDA patients (Fig. 1A) (Saitsu et al., 2013). Polymerase chain reaction (PCR) showed that in mutant mouse two base pairs at the end of exon 6 were removed, resulting in frame-shift mutation and an introduction of a premature stop codon (Fig. 1B). Real-time PCR confirmed that the mRNA of *Wdr45* gene was significantly reduced in mutant mice (referred to as KO thereafter) in the brain (Fig. 1C). Because commercially available antibodies were not able to detect endogenous WDR45 in various tissues in our hands, we used parallel reaction monitoring (PRM), a targeted mass spectrometry approach, to detect a specific WDR45 peptide (YVFTPDGNCR). We were able to detect the WDR45 peptide in WT but not KO mice (Fig. 1D and Supplementary Fig. 1A).

**Figure 1.**
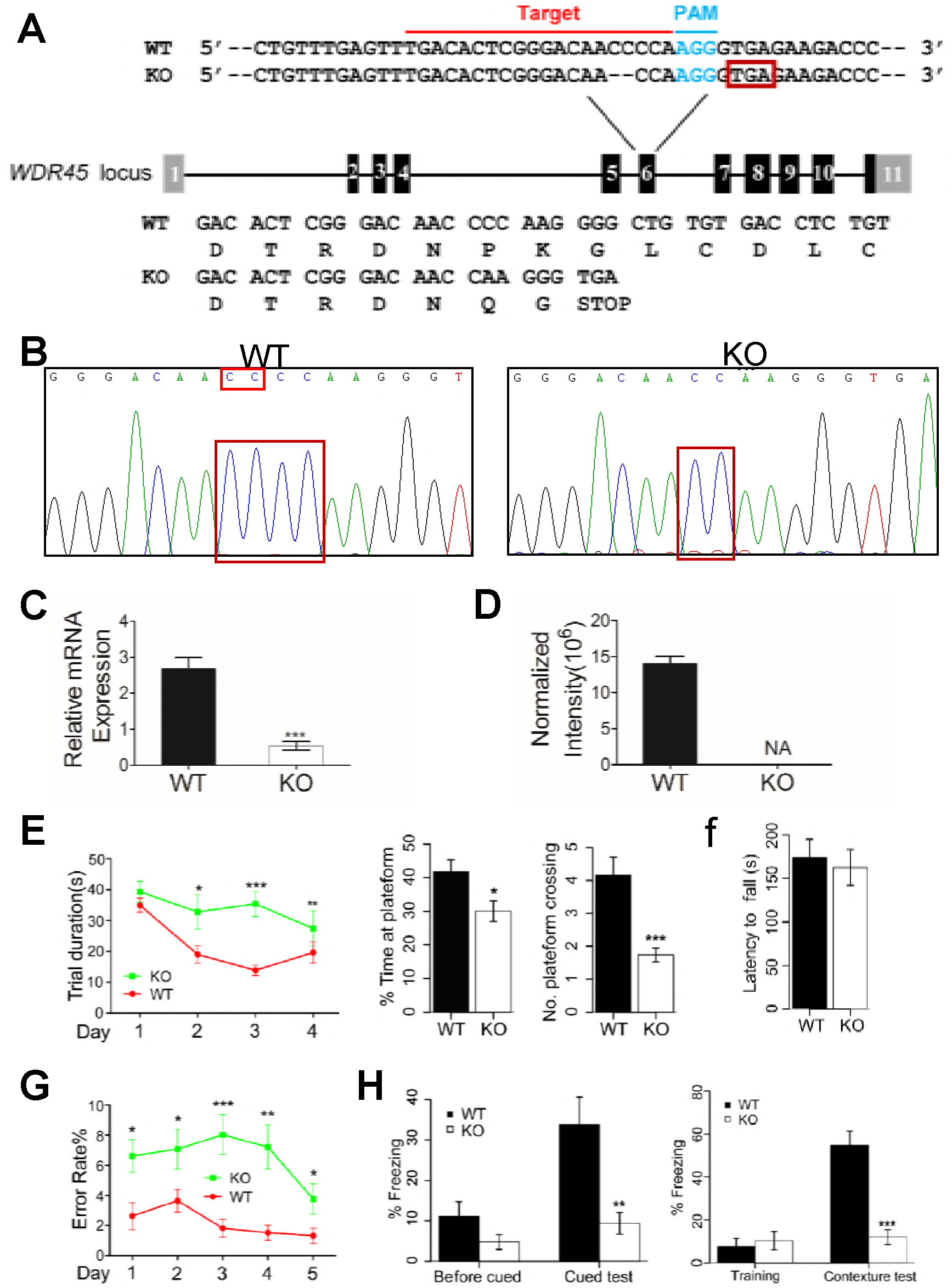
WDR45 knockout (KO) mice show cognitive impairments. (A) Generation of WDR45 KO mouse by deleting a CC dinucleotide at the exon 6 of WDR45 gene. (B) DNA sequencing validation of the deleted dinucleotide in KO mouse. (C) Real time PCR quantification of mRNA expression in WT (n=3) and WDR45 KO (n=3) mice. (D) Protein expression quantified by parallel reaction monitoring (PRM) mass spectrometry of WDR45 target peptide. (E) Morris water maze test of the trial period (left, P<0.01, P=0.006 and P<0.001 on day 2, 3 and 4, respectively. WT: n=12, KO: n=15), time stayed at the platform and platform crossing times (right bar graphs, P= 0.017 and P<0.001 respectively). (F) Rotarod test showing the latency to fall (WT: n=16, KO; n=16). (G) 8-arm maze showing the error rate (*P<0.05, **P<0.001, ***P<0.001. WT: n=15, KO: n=17). (H) Freezing response to the shock in the cued test (left, P=0.0023) and the contexture test (right, P< 0.001), compared to WT mice (n=16 for both WT and KO).

We then performed behavioral experiments for mice at 6 months of age. In the Morris water maze test, we noted significant difference in trial duration between KO and WT mice during the training period. On day 2, 3 and 4 during the training, WT mice spent much less time to find the platform compared to KO mice (Fig. 1E, left, *P*<0.01, *P*=0.006 and *P*<0.001, respectively). During the test, KO mice stayed or crossed the target area with significantly less time compared to WT mice (Fig. 1E, right, *P*= 0.017 and *P*<0.001 respectively). In the 8-arm maze test, the error rate in KO mice was significantly higher than that in WT mice on day 3 (*P*<0.0001), and the error rate significantly declined over time for both KO and WT mice on day 5 (*P*=0.026) (Fig. 1G). The rotarod test did not show significant difference (Fig. 1F). In the feared condition and cued tests, KO mice showed a significantly reduced freezing response to fear in both the contextual and cued stimulation (Fig. 1H). Taken together, our data suggest that at least at the tested age *Wdr45* KO mice showed no sign of impairment in motor coordination. However, they suffer from deficits in spatial learning and conditioning memory.

### *Wdr45* knockout mouse shows mild structural abnormalities in the brain

Since BPAN patients display neurodegeneration and Parkinsonism in adulthood, we examined whether aged mice show signs of neurodegeneration. Haemotoxylin and Eosin (H&E) staining of brain tissues from 16-month-old mice shows that while there appears to be unaltered cell number in prefrontal cortex (PFC) of the KO mice, in substantia niagra (SNR) and caudate putamen (CPu) the cell number is reduced.Particularly in the CPu region the staining of the nuclei becomes weaker, and irregularly shaped neurons can be observed (Fig. 2A).

**Figure 2.**
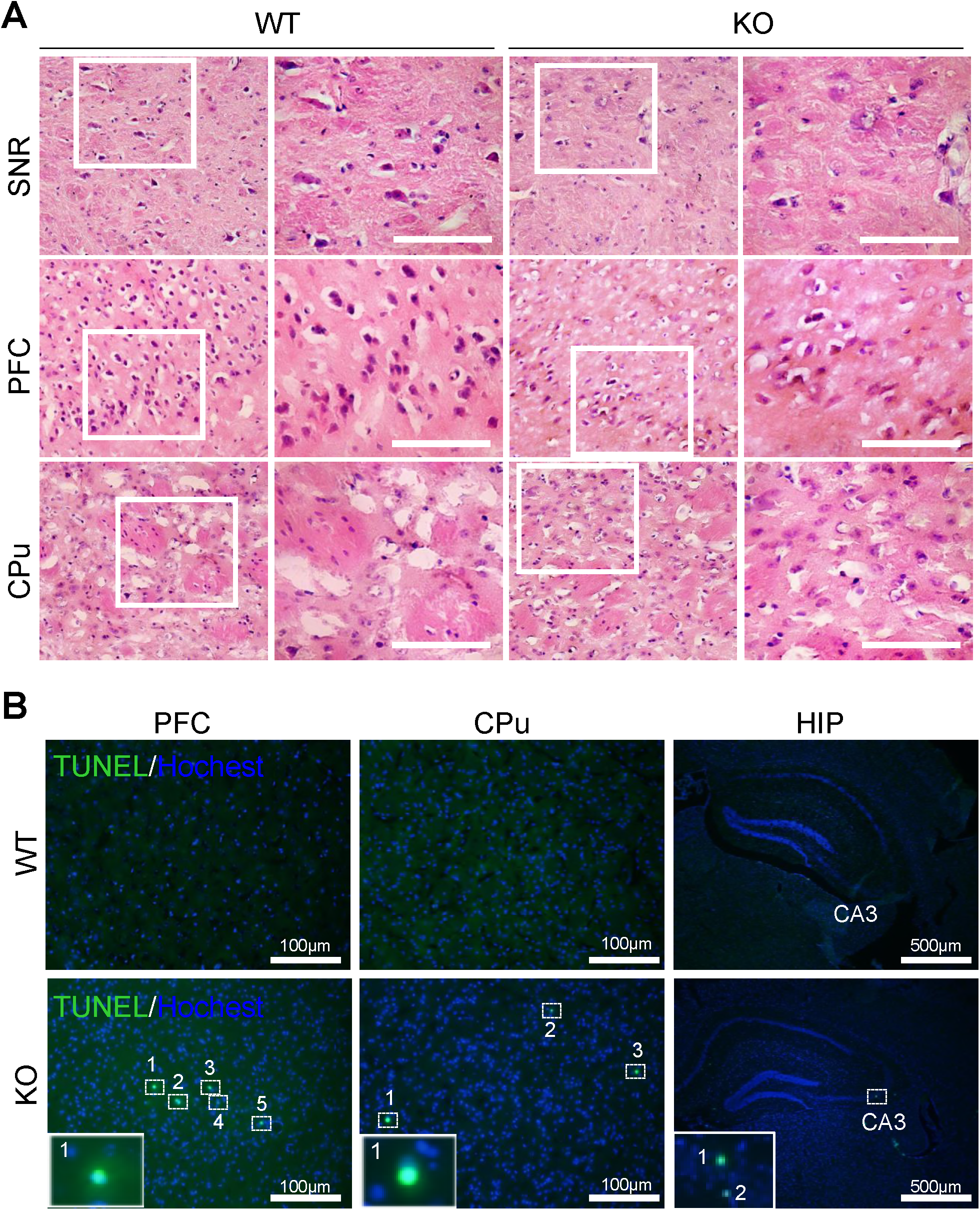
WDR45 KO mice show mild neurodegeneration at old age. (A) H&E staining of brain sections from WT and KO mouse at 16 months of age. Scale bar: 50 μm. (B) TUNEL staining of brain sections from WT and KO mouse at 16 months of age. PFC: prefrontal cortex, CPu: caudate putamen, SNR: substantiao niagra, HIP: hippocampus.

Terminal deoxynucleotidyl transferase (TdT) dUTP Nick-End Labeling (TUNEL) assay on brain regions shows that there are elevated TUNEL labeling in prefrontal cortex (PFC), caudate putamen (CPu), and to a less extent in hippocampus (HIP), in KO mice (Fig. 2B). We also examined whether there are differences in brain iron accumulation, another neuropathological hallmark of BPAN patients. Indeed, we found slightly increased Pearls’ Prussian blue stain of iron in KO mice than WT mice at six month of age. At sixteen month of age the difference became more dramatic (Supplementary Fig. 1B). Thus, the KO mouse we generated shows neurodegeneration at older ages, and mimics at least some pathological features of BPAN patients.

We then used whole-cell patch-clamp recordings to examine the effect of *Wdr45* deletion on the basal synaptic transmission of hippocampal CA1 neurons from 6-month-old mice. We found no significant difference in the frequency of mEPSCs between WT and KO mice (Supplementary Fig. 2A). However, the amplitude of mEPSCs in the CA1 pyramidal neurons was substantially decreased in KO compared to the WT mice, indicating a deficit in post-synaptic function of the hippocampal synapses. In contrast, presynaptic function as assessed by paired-pulse facilitation (PPF) was unaffected (Supplementary Fig. 2B).

Transmission electron microscopy revealed mild defect in synaptic structure (Supplementary Fig. 2C). Although no significant alteration in synapse number, length and width of the post-synaptic density (PSD) was observed in KO brain regions, there was a slight increase in synaptic cleft width in KO brain (Supplementary Fig. 2D). The structural abnormalities in synapse might contribute to the behavioral defects in learning and memory in KO mice.

### Quantitative proteomic analysis of brain regions reveals ER protein accumulation in KO mouse

We dissected five pairs of WT and KO mice brains into pre-frontal cortex (PFC), hippocampus (HIP) and mid-brain (MIB), and performed quantitative tandem mass spectrometry experiments using 10-plex tandem mass tag (10xTMT) labeling (Supplementary Fig. 3A). In total we quantified over 4,000 proteins overlapped in three brain regions, and 67 proteins in PFC, 12 proteins in HIP, and 185 proteins in MIB were significantly changed (*P*<0.05 after Benjamini-Hochberg correction, fold change≥1.5), with PFC and MIB shared 30, and HIP and MIB shared 4 significantly changed proteins (Fig. 3A and B, Supplementary Table 1). The proteins quantified in each individual brain region are listed in Supplemental Tables 2-4. Principal component analysis of the quantitative results shows that the protein expression profiles are well separated between the two genotypes in each brain region (Fig. 3C), indicating that our quantitative mass spectrometry method is able to differentiate mouse brain proteome due to the loss of WDR45. Volcano plots show quantified protein ratios between KO and WT and their *P* values in the three brain regions. Proteins significantly upregulated were labeled red and significantly downregulated labeled green, and we observed more upregulated than downregulated proteins (Fig. 3D). Surprisingly, regardless of well-established role of WDR45 in autophagy, among the 37 quantified proteins intimately participating the autophagy pathway, none of them showed significant change except WDR45 (Supplementary Fig. 3B). To test whether changes in protein level were the results of changes in gene expression, we performed quantitative PCR experiments on 16 significantly changed proteins in the three brain regions and found that with very few exceptions, the mRNA of the majority of the proteins were unaltered (Supplementary Fig. 3C), ruling out the regulation of the protein abundance at the transcription level.

**Table 1.**
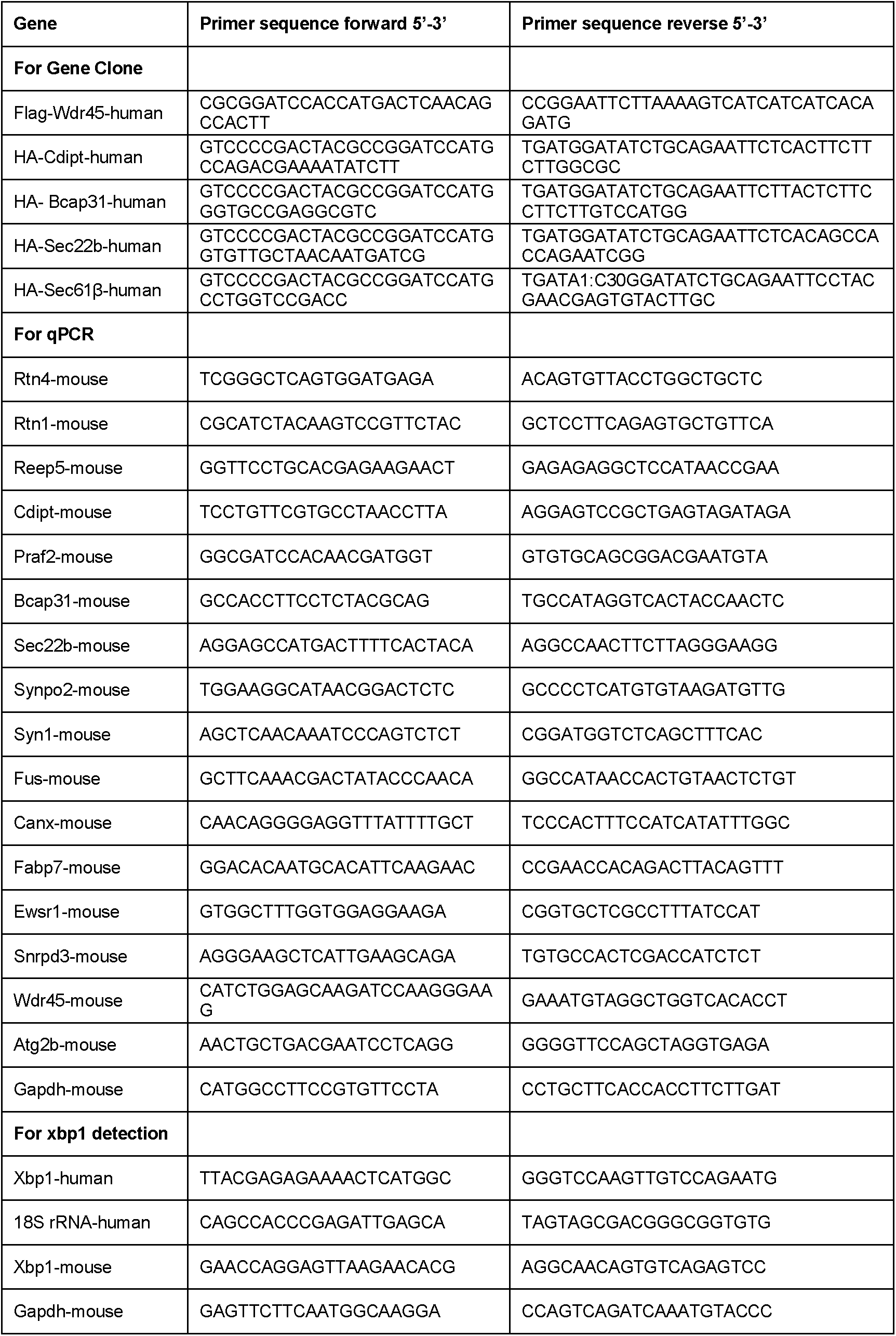
Primer sequences for all the clones and real-636 time PCRs used in the study.

**Figure 3.**
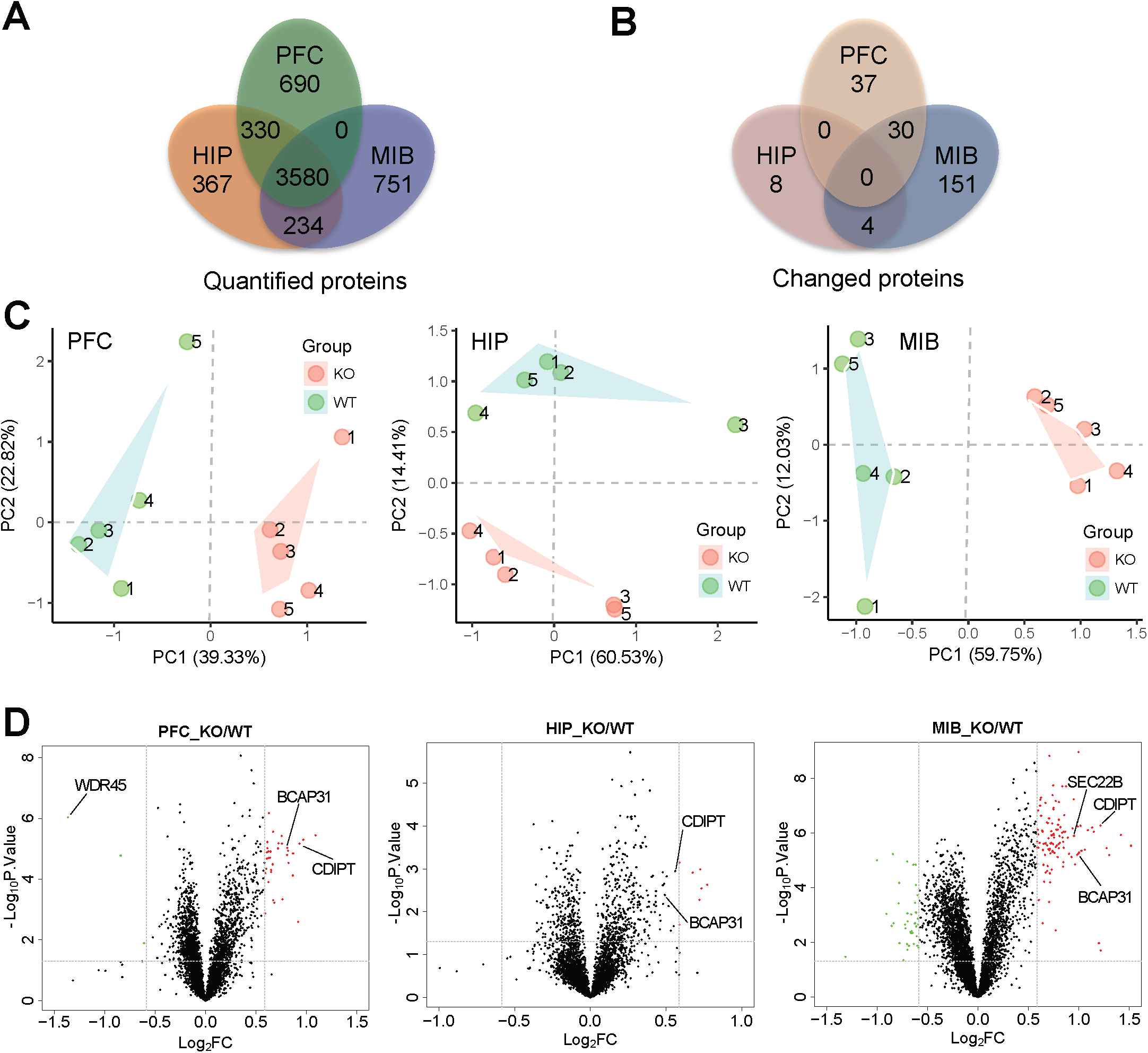
Quantitative proteomic analysis of brain regions from WT and KO mouse. (A) TMT-based mass spectrometry quantified proteins in PFC, HIP, and MIB (n=5). (B) Significantly changed proteins in the three regions (P<0.05 after Benjamini Hochberg correction, and fold change≥1.5). (C) Principal component analysis of proteins quantified in the three brain regions. (D) Volcano plots of quantified proteins in PFC, HIP and MIB, respectively.

Bioinformatics analysis applying Database for Annotation, Visualization and Integrated Discovery (DAVID) (Huang, Sherman et al., 2008, Huang, Sherman et al., 2009) on the changed proteins in MIB and PFC reveals that organization of the ER emerged as the most dramatically enriched gene ontology term, followed by lipid metabolic process and endomembrane system organization (Fig. 4A). Several other cellular compartments including mitochondria and proteasome complex are also among the top ten most enriched terms. Furthermore, weighted gene co-expression network analysis (WGCNA) reveals the accumulation of ER proteins in KO mice (Fig. 4B), and a complex network of ER resident proteins as well as ER-associated proteins shown in the cyan module in Fig. 4C.

**Figure 4.**
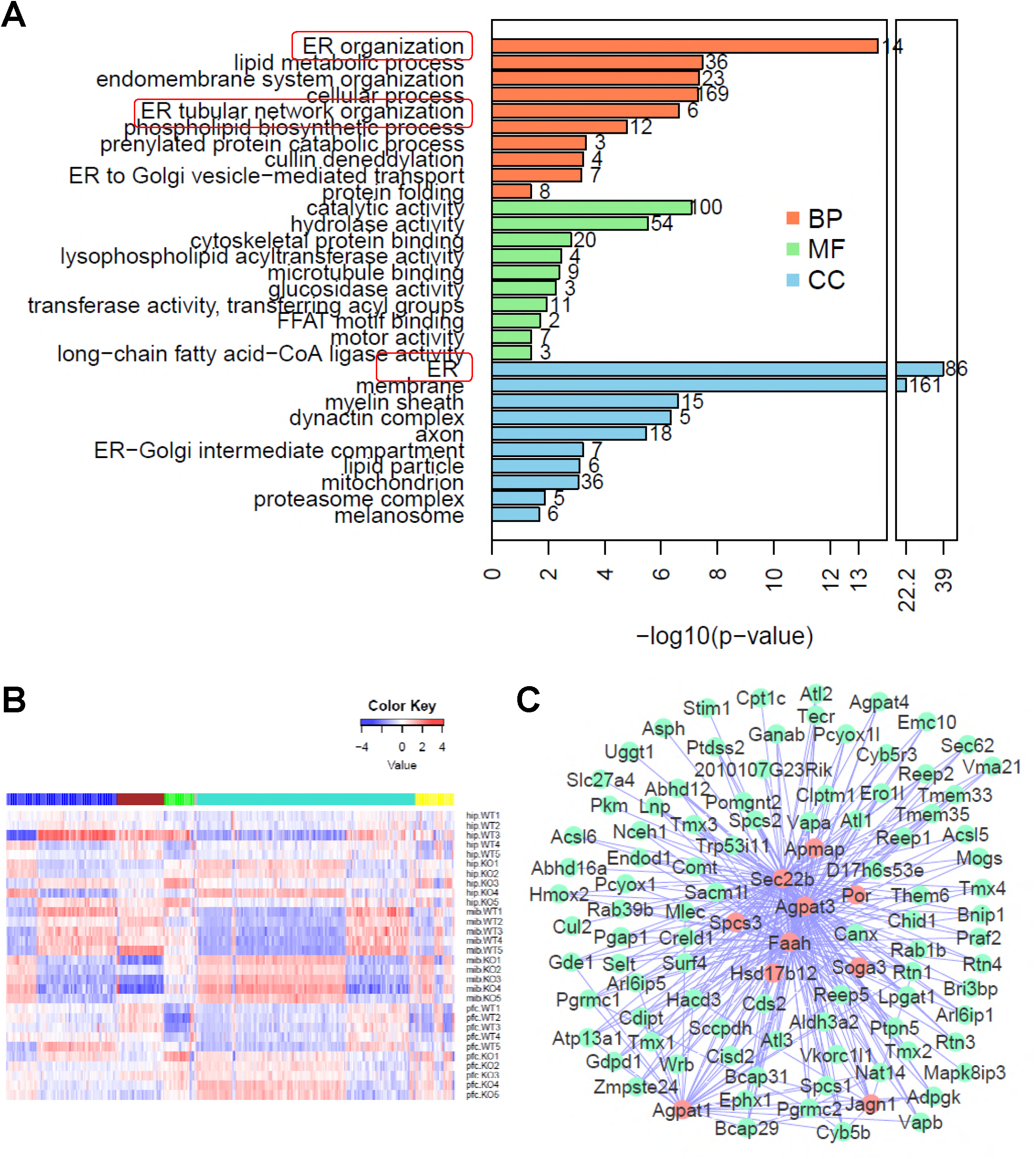
Bioinformatics analysis reveals accumulation of ER proteins in WDR45 mice. (A) Top ten most significantly enriched gene ontology terms in biological process (brown), molecular function (green) and cellular component (blue). (B) WGCNA of proteins quantified in the three brain regions separates the proteins into five distinct co-expressed modules. Color key represent log2-transformed intensity values of each protein. (C) Protein-protein interaction network of the proteins in green module, with red representing hub proteins that have the most network connectivity.

A recent study applied interaction proteomics to identify 321 unique proteins interacting with the six neurodegenerative disease-associated proteins, including Amyloid beta precursor protein (APP), Presenilin-1 (PSEN1), Huntingtin (HTT), Parkin (PARK2), and Ataxin-1 (ATXN1) (Hosp, Vossfeldt et al., 2015). From our quantified union of 5407 proteins from three brain regions, we applied a loosen criteria (P<0.05 and KO/WT ratio > 1.25 or <0.8) to form a list of 678 significantly changed proteins. We mapped these proteins to the 321 proteins and displayed the overlapped 58 proteins with their normalized expression levels in a heat map (Supplementary Fig. 4A). Among the three brain regions, MIB showed the most striking differences between WT and KO mice for the proteins that interact with these disease-associated proteins. The overlap between our dataset and the interaction proteomics data are shown in Supplementary Fig. 4B. Gene ontology analysis captured ubiquitin-proteosome system as the most significantly enriched process, which is in line with the well-established role of protein quality control as an underlying molecular mechanism in neurodegeneration. Of note, endoplasmic reticulum is among the top ten significantly enriched terms (Supplementary Fig. 4C).

### WDR45 regulates target ER proteins via proteasome and lysosome pathways

We then investigated how WDR45 regulates levels of ER proteins. We exogenously expressed Sec22b, Bap31, CDIPT, and Sec61β, four proteins that are either resident ER proteins or ER-associated proteins. When expressed together with WDR45, the levels of all four proteins were significantly reduced (Supplementary Fig. 5A and B). The effect appeared to be specific because co-expression of WDR45 with GFP had no effect comparing to expressing GFP alone (Supplementary Fig. 5C and D).

**Figure 5.**
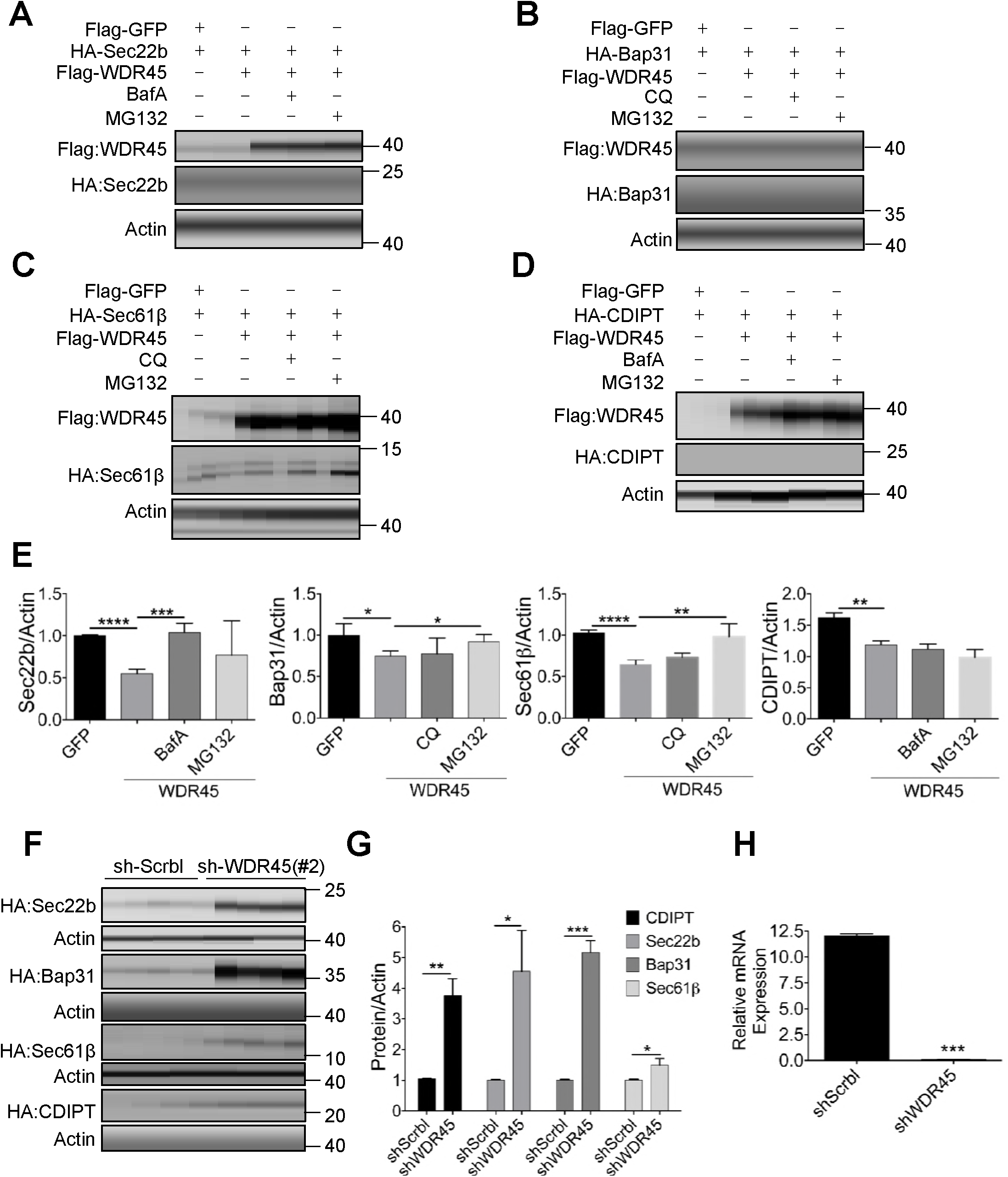
WDR45 mediates degradation of target ER proteins via proteasome and lysosome pathways. (A-D). Co-expression of WDR45 influences the stability of exogenously expressed Sec22b (A), Bap31 (B), Sec61β (C) and CDIPT (D) in Hela cells. Cells were treated with MG132 (20 μM, 12h), BafA (50nM, 8h) and CQ (10 μM, 8h) respectively. (E) Bar graph shows statistics of protein expression levels shown in (A-D). (F) Stability of exogenously expressed Sec22b, Bap31, Sec61β and CDIPT in Hela stable cells with WDR45 knocking down. (G) Bar graph shows statistics of protein expression levels shown in (F). (H) Real-time PCR quantification of WDR45 mRNA in stable cell lines. Data were expressed as means ± SEM (n = 3) and analyzed by two-tailed unpaired t-test. P values: *P<0.05,**P<0.01, ***P<0.001.

To further understand the potential degradation pathway mediated by WDR45, we co-expressed WDR45 and the four proteins in the presence or absence of bafilomycin A (BafA), an autophagy inhibitor that blocks autophagosome-lysosome fusion, or MG132, a proteasome inhibitor. BafA but not MG132 effectively blocked WDR45 mediated degradation of Sec22b (Fig. 5A), while both BafA and MG132 blocked WDR45 mediated degradation of Bap31 (Fig. 5B). For Sec61β, it was MG132 but not BafA that inhibited WDR45 mediated degradation (Fig. 5C), while none of the two inhibitors had any rescue effect on CDIPT (Fig. 5D). As a control, MG132 but not BafA had a mild effect on increasing GFP stability (Supplementary Fig. 5E and F). These different effects suggest that proteins localize at ER at different compartments could be degraded by different mechanisms. However, cells stably expressing shRNA targeting *WDR45* mRNA (sh-WDR45) showed accumulation of all four proteins comparing to cells expressing scrambled (sh-Scrbl) RNA (Fig. 5F and G), similar to the effect observed in KO mouse brain. The knock down effect of *WDR45* mRNA is shown in Fig. 5H.

### Cells with WDR45 deficiency results in ER expansion and increased ER stress

We then investigated whether accumulation of ER proteins correlates to an expanded ER area, a phenomenon that has been recently demonstrated to be the result of reduced selective ER autophagy (Khaminets, Heinrich et al.). Cells expressing shRNA targeting two different *WDR45* mRNA sequences showed significant ER expansion, comparing to cells expressing sh-Scrbl, as imaged by endogenous calreticulin labeling (Fig. 6A and B, *P*<0.001). Similar results were observed by labeling ER using exogenously expressed Bap31, CDIPT, and Sec22b (Supplementary Fig. 6A and B). Furthermore, inducing ER stress using tunicamycin (Tm) results in partial co-localization of WDR45 with calreticulin, while at basal conditions there is no co-localization (Fig. 6C). Transmission electron microscopy shows increased ER engulfment in lysosomes in control cells after ER stress, but the structure is hardly seen in cells with WDR45 deficiency (Fig. 6D). To test whether this effect of selective ER autophagy is organelle specific, we treated cells with a mitochondrial membrane potential uncoupler, carbonyl cyanide m-chlorophenyl hydrazone (CCCP), to induce mitochondrial damage. In cells expressing sh-Scrbl, treatment with CCCP in 24 hours effectively resulted in dramatic loss of parkin staining and shrinkage of mitochondrial area, suggesting parkin-mediated mitophagy. However, in cells expressing sh-WDR45, the mitochondrial area appeared significantly larger than control cells (Supplementary Fig. 6C). Thus, we demonstrated that a defect in macroautophagy resulted in defects in selective autophagy of ER and mitochondrion, two independent organelles.

**Figure 6.**
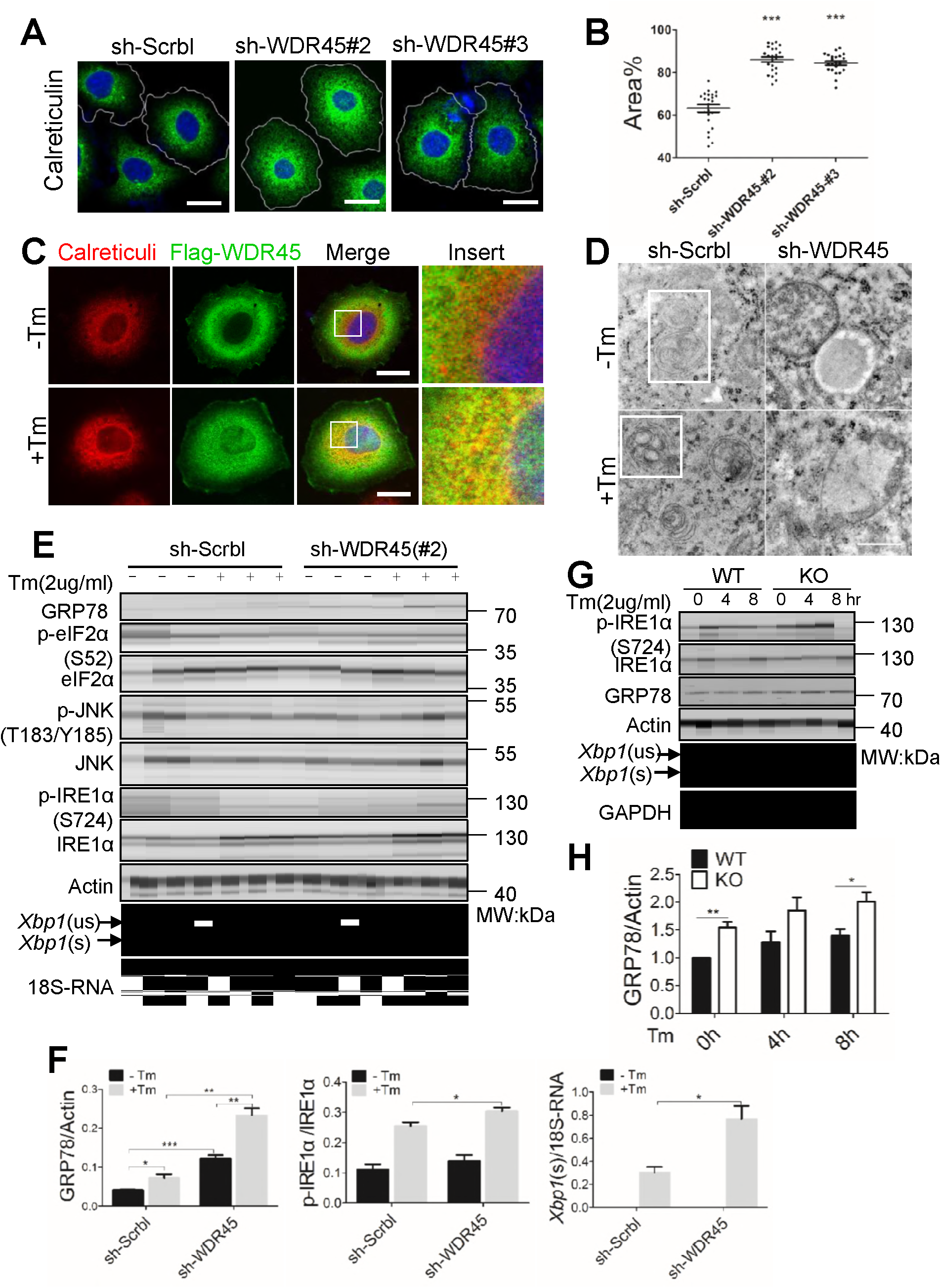
WDR45 deficient cells display increased ER stress and ER area. (A) Representative confocal images of immunofluorescent labeled ER resident protein calreticulin in Hela cells. Scale bars, 20 um.(B) Quantification of ER area, ***P<0.001, unpaired t test, n=30. (C) Co-staining of Flag-WDR45 with calreticulin in Hela cells with or without Tm treatment (2 μg/ml, 8 hrs). Scale bars, 20 μm. (D) Representative TEM images capturing ER engulfment in lysosomes in Hela cells with or without Tm treatment (2 μg/ml, 8 hrs). Scale bar, 500 nm. (E and F) Hela cells, (G and H), primary cortical neurons were treated with Tm, immunoblotted with antibodies in the ER stress pathways. Data were expressed as means ± SEM (n = 3) and analyzed by two-tailed unpaired t-test. **P<0.05, **P<0.01, ***P<0.001.

What could be the physiological upstream trigger for compromised organelle integrity? A well-established mechanism to handle ER stress accumulated over time is the unfolded protein response (UPR) (Frakes & Dillin, 2017, Hetz & Saxena, 2017). We therefore examined this pathway and found that the ER chaperon GRP78 was significantly upregulated in cells expressing sh-WDR45 comparing to control cells under basal condition. Inducing ER stress using Tm further promoted GRP78 expression (Fig. 6e and f, and Supplementary Fig. 6E). Out of the three known UPR pathways (Frakes & Dillin, 2017, Hetz & Saxena, 2017), we found over activation of the IRE1α pathway, because there was statistically significant upregulation of phosphorylated IRE1α at serine 724 in cells expressing sh-WDR45. In addition, we observed an increased selective splicing product of Xbp1 (Fig. 6E and F). In contrast, we found no evidence of significant change in eIF2 phosphorylation, an indication of PERK activation (Fig. 6E). In primary neurons from KO mouse, we also found increased GRP78 expression, IRE1α activation and increased selective splicing of Xbp1 (Fig. 6G and H).

### WDR45 deficiency leads to an increased cell death after ER stress

If an abnormally increased ER stress cannot be properly handled by UPR or selective autophagy, it could eventually leads to apoptosis (Hetz & Saxena, 2017). Cell viability assay shows that Tm treatment resulted in significantly reduced viability in cells expressing sh-WDR45 than control cells (Fig. 7A), the difference in cell viability is further supported by increased caspase-3 cleavage (Fig. 7B and C). In primary neurons with WDR45 deficiency, there are increased level of the ER marker calreticulin, accumulation of the autophagy receptor p62, and reduced LC3II/LC3I levels (Fig. 7D, Supplementary Fig. 7A). The defect in autophagy as marked by p62 accumulation and reduced LC3II/LC3I levels, increase ER stress as marked by GRP78 and activation of the IREα pathway, and finally increase apoptosis were all recapitulated in brain tissues by immunoblot and immunohistochemistry (Fig. 7E and F, Supplementary Fig. 7C-E). These defects were not observed in 1-month-old WDR45 KO mouse (Supplementary Fig.7 B), which indicates WDR45 has negligible influence on the early stage of development.

**Figure 7.**
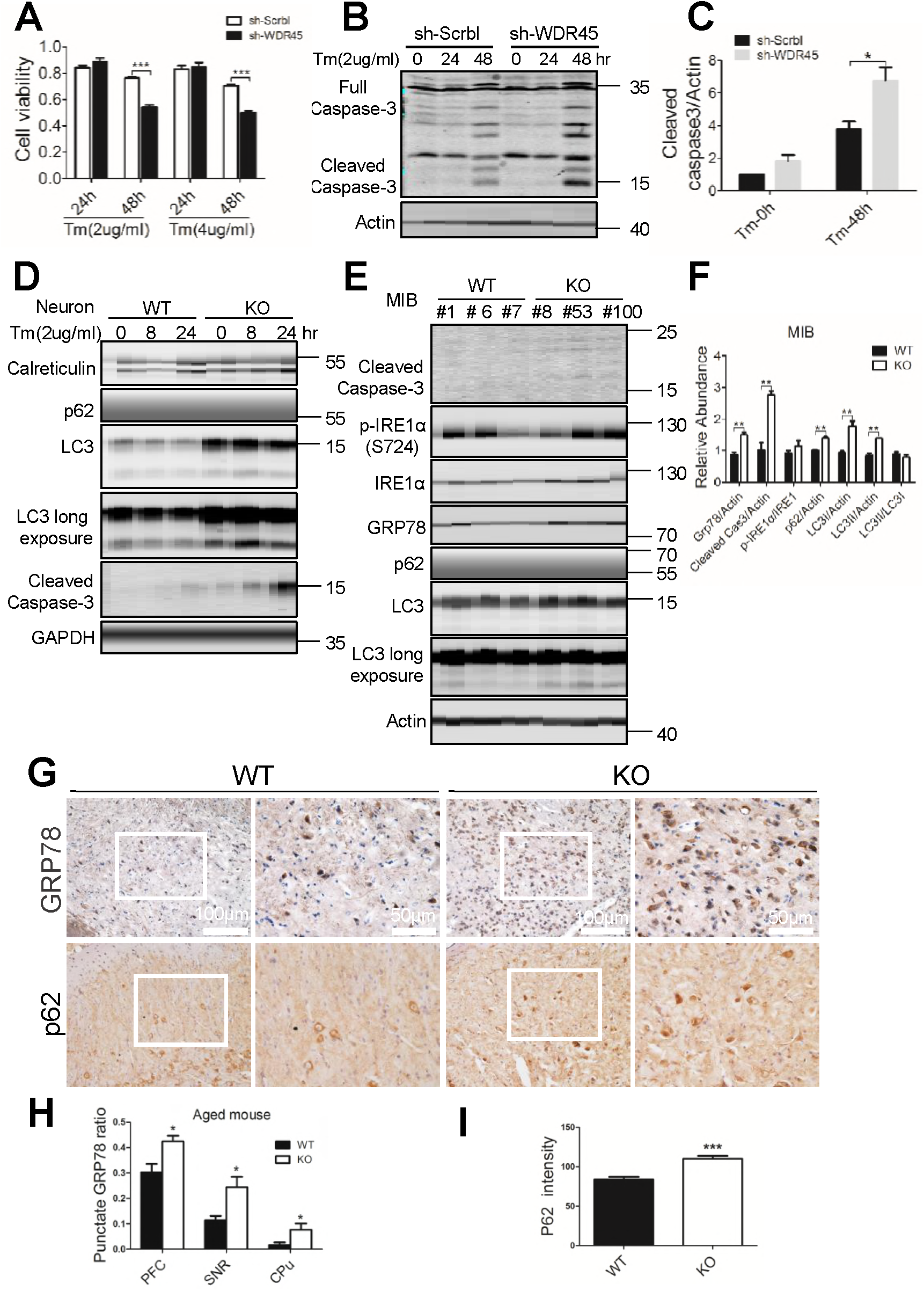
Loss of WDR45 results in an increased cell death associated with ER stress. (A) Cell viability was measured in sh-Scrbl and sh-WDR45 Hela cells after Tm treatment. Data were expressed as means ± SEM (n = 6) and analyzed by two-tailed unpaired t-test. ***P < 0.001. (B) Western blot analysis of caspase-3 cleavage after Tm treatment in Hela cells. (C) Statistical analysis of (B). (D) Western blot analysis of caspase-3 and autophagy proteins after Tm treatment in WT and KO primary neurons. (E) Western blot analysis of caspase-3, ER stress markers and autophagy proteins in midbrain from WT and KO mouse at 18-20 months of age. (F) Statistical analysis of (E). (G) Immunohistochemistry of GRP78 and p62 in substantia niagra. (H and I) statistical analysis of GRP78 (16 months) and p62 (6 months), respectively. Data were expressed as means ± SEM (n = 3) and analyzed by two-tailed unpaired t-test. *P<0.05, **P<0.01, ***P<0.001.

### Activation of autophagy or inhibition of ER stress rescues apoptosis

We further asked whether pharmacological rescue of apoptosis could be achieved by modulating the levels of autophagy and ER stress. We used rapamycin, an inhibitor of mTOR signaling that could potentially relieve the inhibition of autophagy by mTOR (Kim & Guan, 2015), and tauroursodeoxycholic acid (TUDCA), an ER stress inhibitor (Keestra-Gounder, Byndloss et al., 2016). Either rapamycin or TUDCA reduced accumulation of exogenously expressed ER proteins, Sec22b and Sec61β, in WDR45-deficient cells (Fig. 8A). Aggravated ER stress after Tm induction in WDR45-deficient cells was reduced by rapamycin or TUDCA (Fig. 8B). Increased caspase-3 cleavage in WDR45-deficient cells was also reduced by rapamycin or TUDCA (Fig. 8C). Flow cytometry labeling of cell surface markers Annexin V and DNA marker 7-ADD for apoptosis also demonstrated that Tm induced apoptosis was alleviated by treatment with either rapamycin or TUDCA (Fig. 8D). In cultured primary WT but not KO neurons, Tm strongly induced autophagic activity as demonstrated by increased LC3II/I ratio, indicating a defect in autophagy in KO neurons. Treatment with rapamycin but not TUDCA mildly but significantly increased LC3II/I ratio, suggesting a partial restore of autophagic activity (Fig. 8E). Finally, rapamycin significantly reduced capase-3 cleavage in KO neurons, although TUDCA had similar trend of reduction but did not reach statistical significance (Fig. 8E). Together, our data provide an integral link between a defect in autophagy due to WDR45 deficiency, ER stress, and apoptosis as an underlying mechanism of β-propeller protein associated neurodegeneration.

**Figure 8.**
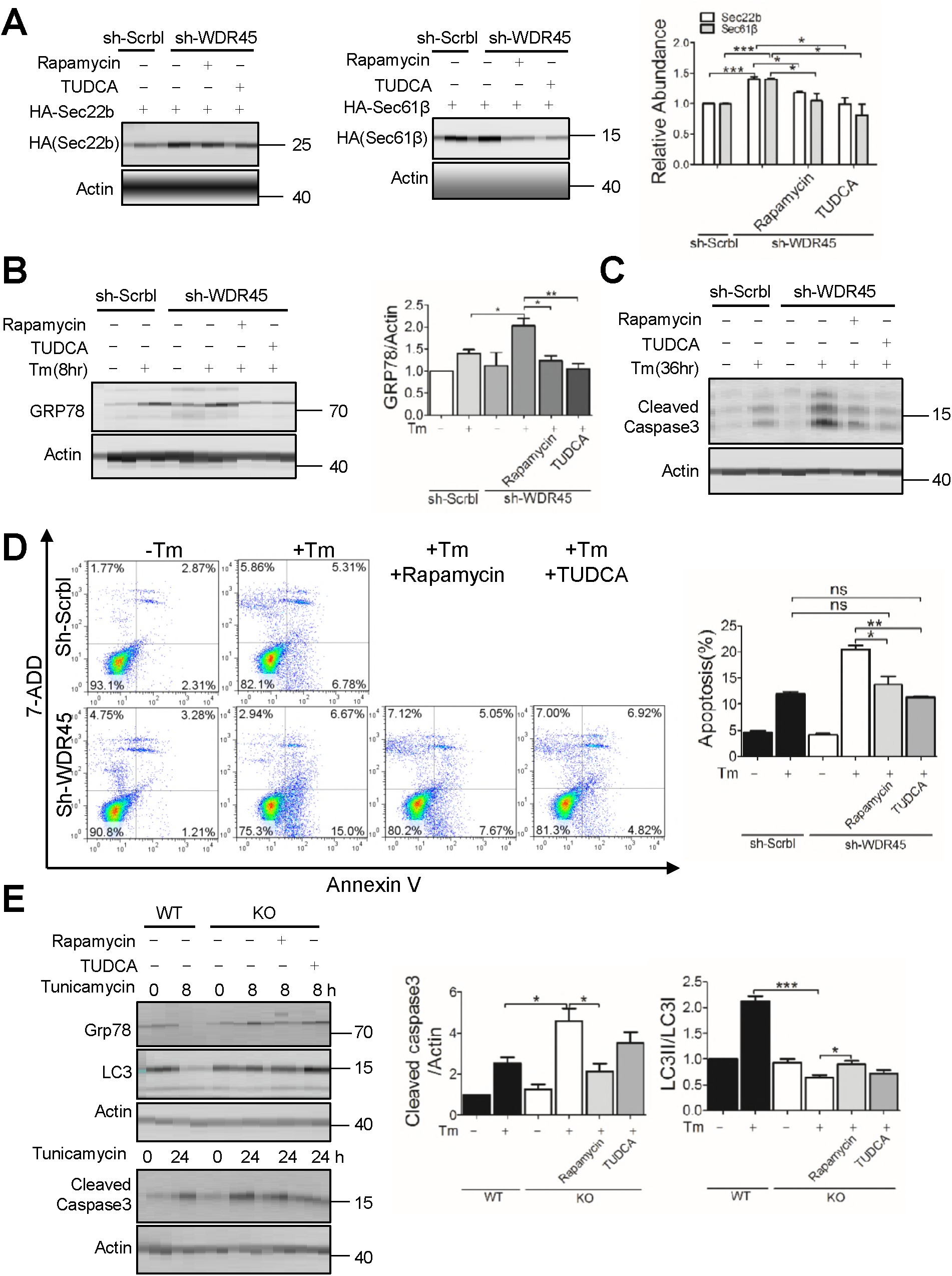
Inhibition of mTOR or UPR rescues abnormal protein processing and increased cell death after loss of WDR45. (A) Western blot analysis of protein stability after rapamycin (100 nM, 24hr) or TUDCA (100 μM, 24hr) treatment in WDR45 deficient cells. (B) Effect of rapamycin or TUDCA on ER stress marker GRP78 in WDR45 deficient Hela cells after inducing ER stress. (C) Effects of rapamycin (100 nM, 36h) or TUDCA (100 μM, 36hr) on caspase-3 levels in WDR45 deficient Hela cells after inducing ER stress. (D) Flow cytometry analysis of apoptosis in WDR45 deficient Hela cells after inducing ER stress (2 μg/ml Tm, 36hr). (E) Effects of rapamycin or TUDCA on Grp78 accumulation (Tm, for 8hr), caspase-3 cleavage (Tm, 24hr), and LC3II production, in neurons after inducing ER stress. Data were expressed as means ± SEM (n = 3) and analyzed by two-tailed unpaired t-test. *P<0.05, **P<0.01, *** P<0.001.

## Discussion

Although genetic bases of β-propeller protein-associated neurodegeneration has been well established, the molecular and cellular underpinnings of the disease process are currently not yet known. Our study filled this gap by providing compelling evidence showing that a deficit in autophagy due to *Wdr45* mutation leads to improperly handled organelle stress, particularly ER stress, which is accumulated over time of the mouse life span. Accumulation of ER stress triggers UPR through IRE1α and possibly other kinase signaling pathways, and eventually leads to neuronal apoptosis, a hallmark of neurodegeneration. Futhermore, we show that suppression of ER stress, or activation of autophagy through inhibition of mTOR pathway, rescues neuronal death.

A previous study generated neuronal-specific knockout of *Wdr45* in mouse and observed poor motor coordination and deficits in learning and memory at 9-12 months of age, and axonal swelling at 11-13 months (Zhao et al., 2015). To model *de novo* mutation of *WDR45* in BPAN patients, we generated a germ line mutation in mouse copying a key mutation found in patients, and resulted in a knockout genotype. Behaviorally, although we found no significant alteration in motor coordination, we found significant impairment in both spatial and contextual memory as early as 6 months of age (Fig. 1E-H). Similarly, our KO mice show abnormalities in basal synaptic transmission and mild structural defects in the synapse (Supplementary Fig. 2). Thus, our germ line mutation mouse shares some features with that of conditional knockout mouse. In addition, our KO mice also display distinct pathological features such as brain ion accumulation at basal ganglia and increased neuronal death at older age (Fig. 2 and Supplementary Fig. 2), largely in line with pathological features found in BPAN patients. However, although behavioral and electrophysiological experiments show early impairments in hippocampal function, we observed very mild neurodegenration in hippocampus even at old ages, suggesting that other cellular pathways exist to account for the synaptic and behavioral abnormalities found in this brain region.

Our study provide direct evidence showing that the defect of a protein associated with macroautophagy can results in defects in selective autophagy in organelles. To understand the selective vulnerability of ER to WDR45 mutation, we have affinity purified overexpressed WDR45 complex from cells after induction of ER stress. Mass spectrometry results showed that only ATG2B was the most prominent protein partners of WDR45, confirming previous reports (Behrends, Sowa et al., 2010). We also attempted to test whether known ER selective autophagy receptors interacted with WDR45, including FAM134B, Sec62, and RTN3 (Fumagalli, Noack et al., 2016, Grumati, Morozzi et al., 2017, Khaminets et al., 2015), however none of which showed clear interaction. It is thus possible that mutation of WDR45 results in low autophagic activity in general, which affects macro-as well as selective autophagy. Accumulation of misfolded proteins in neurons induces ER stress, which can be handled properly in wild type neurons. Selective ER autophagy maybe achieved through elevated expression of ER specific receptors including FAM134b or Sec62 (Fumagalli et al., 2016, Khaminets et al., 2015) to provide a destruction signal. Neurons with lower autophagic activity due to WDR45 mutation amplifies ER stress over time, through a compromised ability to degrade damaged ER and protein aggregates, even though selective ER autophagy signals are intact. Our parallel study of mitochondrial also shows a compromised mitophagy in WDR45-deficient cells, further support this notion that selectivity of organelle autophagy maybe achieved through selective stress of the organelle.

ER stress has emerged as a common cellular pathway underlying neurodegeneration. As the results of mishandled ER stress, protein aggregates in various brain regions have been found in a number of neurodegenerative disorders, including Alzheimer’s disease, Parkinson’s disease, frontotemporal dementia, and Amyotrophic lateral sclearosis (Hetz & Saxena, 2017). Through an unbiased quantitative proteomic analysis of mouse brain due to WDR45 knocking out, we uncovered accumulation of large number of ER proteins. Our follow-up functional analysis indeed revealed abnormal UPR pathway that is presumably upregulated to handle increased ER stress. Thus, our study adds another neurodegenerative disorder whose mechanistic onset is at least partly due to increased ER stress, sustained activation of UPR pathway.

Intriguingly, either inhibition of mTOR signaling or inhibition of ER stress is able to rescue apoptosis in WDR45-deficient cells and neurons. Although inhibition of mTOR could have a wide range of effects on catabolic processes, our data did show an accompanying increase of LC3II/LC3I ratio, a maker of increased autophagy. TUDCA is known to exert several effects that lead to neuroprotection, including inhibition of mitochondrial pathway of cell death, reducing oxidative stress, and reducing ER stress (Özcan, Yilmaz et al., 2006); it has been applied in animal models of Alzheimer’s, Parkinson’s, and Huntington’s disease and show at least some positive effects against neuronal death (Castro-Caldas, Carvalho et al., 2012, Keene, Rodrigues et al., 2002, Nunes, Amaral et al., 2012). It is thus not entirely surprising that TUDCA inhibit apoptosis in WDR45-deficient cells to the extent similar to mTOR inhibition. In neurons, however, only mTOR inhibition significantly reduced apoptosis, further highlight the importance of autophagy in maintaining protein as well as organelle homeostasis in neurons. It is worth noticing that neither inhibition of mTOR nor inhibition of ER stress rescued the ER expansion phenotype in WDR45-deficient cells, at least in the experimental setting similar to rescue of apoptosis. We postulate that the level of ER autophagy induced by rapamycin or TUDCA may not be sufficient to manage the complex ER network. Alternatively, WDR45 may be involved in other pathway to control ER quality. Nevertheless, our study suggests a unified principle underlying neurodegeneration regardless of a variety of etiological origins, and highlight potential cellular pathways that could be modulated to reduce neurodegeneration in BPAN patients.

## Methods

### Generation of *Wdr45* knockout mouse

The mouse was generated by co-injection of Cas9 mRNA and sgRNA (target sequence: ACACTCGGGACAACCCCA), as described previously (Shao, Guan et al., 2014). Briefly, superovulated female C57BL/6J mice were mated to male C57BL/6J mice, and embryos were collected from oviducts. Cas9 mRNA (100 ng/μl) and sgRNA (50 ng/μl) were co-injected into the pronuclei of one-cell embryos. The injected embryos were cultured in KSOM medium overnight before they were transplanted into pseudopregnant mice. One week after birth, genomic DNA from the toes or tail of the newborn F0 mice was extracted for sequencing. Mice were housed in standard cages in a specific pathogen-free facility on a 12-h light/dark cycle.

### Motor coordination

The motor coordination was measured as described previously with some modifications. After trained on the rolling rod at 5 rpm, 5-15 rpm and 8-25 rpm for 5 min for three days, mice were tested for their ability to remain on the rotarod at 8-25 rpm. The time when the mice fell from the rod was recorded with a maximum of 5 min.

### Morris water maze

A round tub was filled with opaque water (20°C) to a depth of 25 cm. A platform was located in the center of one quadrant. In the learning test, the mouse was placed in the maze starting from one of four predetermined spots and was given 60 seconds to find the fixed platform. The time taken to reach the platform (escaping latency) was recorded by a video tracking system. Each mouse received 4 trials per day for 4 consecutive days. On the 5th day, the probe test was performed without the platform. The mouse was allowed to search for 60 seconds and the percentage of time spent in the target quadrant was recorded to monitor spatial memory ability.

### Eight-arm maze

Mice were individually placed into the center of a symmetrical 8-arm maze and allowed to freely enter the arms during a 10-minutes session. The series of arm entries was recorded visually. The percentage of alternation was calculated as the ratio of correct alternations (successive entry into the 8 arms on overlapping triplet sets) to total alternations.

### Feared conditioning

The fear conditioning test was performed using the Startle and Fear combined system (Panlab,Harvard Apparatus). For training, after 3 minutes acclimation, mice were placed in a chamber and an 80 dB tone was delivered for 30 seconds as a conditioned stimulus. During the last 2 seconds, a foot shock of 0.4 mA was delivered through a shock generator. Animal movement was recorded through a high-sensitivity weight transducer system by PACKWIN software. On the second day, for the contextual test, mice were placed in the chamber and the freezing response was measured for 3 minutes without the conditioned stimulus. For the cued test, the freezing response was measured with the conditioned stimulus.

### Slice electrophysiology

Experiments were performed on hippocampal slices from WT and KO mice as described previously (Zhou, Liu et al., 2018). Briefly, after decapitation, the mouse brain was quickly removed and placed in well-oxygenated (95% O_2_/5% CO_2_) ice-cold artificial cerebrospinal fluid (aCSF) containing (in mM): 125 NaCl, 2.5 KCl, 12.5 D-glucose, 1 MgCl_2_, 2 CaCl_2_, 1.25 NaH_2_PO_4_ and 25 NaHCO_3_ (pH 7.35–7.45). Coronal hippocampal slices (300 mm thick) were obtained with a vibratome (Leica VT 1000S) and incubated at 30 ± 1°C in oxygenated ACSF for at least 1 h before being transferred to a recording chamber placed on the stage of an Olympus microscope (BX51WI). The slices were continuously perfused with oxygenated aCSF at room temperature during all electrophysiological studies. Hippocampal CA1 neurons were visualized using an infrared differential interference contrast microscope. Synaptic responses were recorded with an Axon 200B amplifier (Molecular Devices), and the stimulations were delivered with a bipolar tungsten-stimulating electrode placed over Schaffer collateral fibers between 250 and 300 μm from the recorded cells in the CA1 region. Pyramidal neurons were identified based on their ability to exhibit spike frequency adaptation in response to prolonged depolarizing current injection. AMPAR-mediated EPSCs were induced by repetitive stimulations at 0.03 Hz, with the neuron being voltage-clamped at –70mV except where indicated otherwise. The recording pipettes (4–7 MΩ) were filled with a solution containing (in mM): 145 potassium gluconate, 5 NaCl, 10 HEPES, 2 MgATP, 0.1 Na_3_GTP, 0.2 EGTAand 1 MgCl_2_ (280–300 mOsm, pH 7.2 with KOH). Paired-pulse facilitation was induced by delivering two consecutive pulses with a 25-, 50-, 75-, 100-, or 200ms inter pulse interval and calculated as the ratios of the second response peak values over the first response peak values. The miniature EPSCs of CA1 pyramidal neurons in the hippocampus were obtained at -70 mV in the presence of tetrodotoxin (0.5 mM) and picrotoxin (100 mM) without stimulation and analysed using MiniAnalysis (Synaptosoft). In all of the whole-cell recordings, cell series resistance was monitored throughout experiments and the experiment was excluded from analysis if the resistance changed by more than 20%.

### Chemical reagents, antibodies and plasmids

Rapamyicin (Sigma-Aldrich, #V900930), TUDCA (Tauroursodeoxycholic acid, Shang Hai Yuan Ye, B20921, CHN), Tunicamycin (Cell Signaling Technology, #12819), BafA (Sigma-Aldrich, #B1793), and MG132 (Selleck, #S2619) were used at final concentrations of 100 nM, 100 μM, 2-4 μg/ml, 50 nM and 20 μM, respectively.

The following antibodies were used in the western blot experiments: rabbit anti-HA (1:1000, Cell Signaling Technology, #12698), mouse anti-β-actin (1:5000, YEASEN, #30101-ES10, CHN), mouse anti-Flag (1:5000, Sigma-Aldrich, #1804), rabbit anti-Calreticulin (1:1000, Abcam, #ab92516), rabbit anti-p62 (1:1000, Sigma-Aldrich, #B0062), rabbit anti-GAPDH (1:5000, Abways, #AB0037, CHN), rabbit anti-cleaved caspase3 (1:1000, CST, #9664), rabbit anti-Tomm20 (1:500 for IF, Santa Cruz Biotechnology, sc11415), rabbit anti-LC3 (1:1000, Sigma-Aldrich, 2775S), and rabbit anti-GRP78 (1:1000, Abways, #CY5166, CHN). Rabbit anti-eIF2α, rabbit anti-IRE1α, rabbit anti-phospho-IRE1α (S724), rabbit anti-JNK, rabbit anti-phospho-JNK (T183/Y185), rabbit anti-phospho-eIF2α (S52) and rabbit anti-Calnexin are all kind gifts from Dr. Liu Yong (Institute of Neuroscience, SIBS, CAS, CHN).

All plasmids were verified by DNA sequencing. The plasmids flag-WDR45, flag-GFP, HA-sec22b, HA-sec61β, HA-Bap31, HA-CDIPT were cloned from pcDNA3.1(+)-flag or HA-CMV vector. All primer sequences are listed in Table 1.

### shRNA knock down

Annealed oligonucleotides were cloned into pLKO.1puro using AgeI and EcoRI cloning sites. The following pairs of oligonucleotides were used to target WDR45: #2: CCGGCAAGATCGTGATCGTGCTGAACTCGAGTTCAGCACGATCACGATCTTGTTTTTTG and AATTCAAAAAACAAGATCGTGATCGTGCTGAACTCGAGTTCAGCACGATCACGATCTTG, #3: CCGGCAAGAACGTCAACTCTGTCATCTCGAGATGACAGAGTTGACGTTCTTGTTTTTTG and AATTCAA AAAACAAGAACGTCAACTCTGTCATCTCGAGATGACAGAGTTGACGTTCTTG. Hela cells stably expressing shRNA oligonucleotides were generated by lentiviral infection as described previously (Popovic, Akutsu et al., 2012).

### RNA extraction, reatl-time PCR

Total RNA was extracted using TRIzol (Gene solution, #B140101) and cDNA was synthesized by Prime ScriptIIFirst-Strand cDNA synthesis Kit (Invitrogen, #11752050) according to the manufacturer’s instructions. Quantitative PCR was performed on a real-time PCR system using SYBR Premix (Promega, #A6001). Primer sequences are listed in Table 1.

### Cell culture and transient transfection

HEK293T and HeLa cells were maintained in DMEM (Invitrogen, #11965) supplemented with 10% FBS (GEMINI) and 50 mg/ml penicillin/streptomycin at 37°C in 5% CO_2_. Cells grown at low confluence in 6 cm, 12-or 6-well tissue culture plates were transfected with 4 μg, 1 μg or 2 μg of plasmid DNA, respectively, using the polyetherimide reagent (Polyseciences, #690049). Experiments were performed 24-36 h after transfection.

Primary cortical neurons were harvested from embryonic day 18 mouse pups. Cerebral cortices were collected and dissociated by incubation in Papain (sigma, #1495005) for 30 min at 37°C. After termination with FBS, cells were centrifuged at 800rpm for 2 min, and plated with MEM media (Gibco, #41090093) on dishes pre-coated with poly-d-lysine (100 mg/ml, Sigma, P1524). After incubation at 37°C for at least 4 h, the plating medium was changed to Neurobasal medium containing B27 (Invitrogen, #17504044), penicillin/streptomycin (Gibco, #15140163) and Glutamine (Invitrogen, #35050061) and 5% FBS (HyClone Laboratories, #SH3008403). Half the volume of culture medium was changed every 3 days.

### Immunoblotting

Cells were lysed with RIPA buffer (50 mM Tris-HCl, pH 7.4, 150 mM NaCl, 1 mM EDTA, 0.1% SDS, 1% NP40) supplemented with protease inhibitor (Thermo Scientific, #88666), sonicated and incubated on ice for 30 min. Homogenates were centrifuged at 13,000 rpm for 20 min at 4°C. Supernatant fractions were collected and protein concentrations were determined by BSA (Thermo Scientific, #23225). Equal amounts (20 to 50 μg) of proteins were subjected to SDS-PAGE electrophoresis, transferred to a nitrocellulose filte membrane and then blocked with 5% nonfat milk for 1 h at room temperature, membranes were incubated with primary antibodies at 4°C overnight. After washing with TBST (0.1%Tween in TBS) 3 times, the membranes were incubated with secondary antibodies for 1 h at room temperature.

### Immunofluorescence (IF) microscopy

Hela cells plated on glass coverslips were treated with indicated drugs. Cells were then washed twice in PBS and fixed at room temperature for 30 min with 4% paraformaldehyde diluted in PBS, permeabilized with 1% Triton X-100 on ice for 10 min, blocked with 5% BSA in 37°C incubator for 60 min and incubated with primary antibodies overnight. The coverslips were washed 3 times with PBST, followed by incubation with a secondary antibody (Life Technology, Alexa Fluor 405-blue, 546-red, 488-green). Confocal pictures were acquired using a Leica TCS SP5 microscope with a 63×/1.4 N.A. objective.

### Immunohistochemistry

Mice were trans-cardially perfused, and cerebral tissues were dissected and postfixed in 4% PFA for 1 day, dehydrated by 20% and 30% sucrose PBS buffer, sectioned at 15um by Leica CM1860. Slices were stained by H&E staining (Sigma-Aldrich) and Perls’ staining (SenBeijia, #SBJ0413, CHN) was performed according to manufacturer instructions.

For immunohistochemistry, slices were stained with a DAB kit (NEOBIOSCIENCE, #ENS004, CHN), briefly, slices incubated with 0.01 M sodium citrate (pH=6.02) at 95°C for 10 min, and inactivated peroxidase with 3% H_2_O_2_,blocked with goat serum and incubated with indicated primary antibodies at room temperature for 2 hours. After washing, sections were incubated with biotin-IgG followed by Streptomycin-HRP incubation for 15 min at room temperature. Finally, sections were counterstained with DAB, and stained DNA with hematoxylin. Pictures were acquired using an Olympus-KCC-REM microscope.

### Transmission electron microscopy

Mice were deeply anesthetized with an intraperitoneal injection of a mixture of 4% chloral hydrate (about 0.01ml/g mouse body weight, respectively), then perfused with 0.1 M phosphate buffer at flow rate 6.5 r/min for 5 min, then perfused with 4% PFA solution in 0.1 M phosphate buffer for 15 min. The whole brain was dissected free with the BRAIN BLOCKER that designed to reproduce the plane of section of the Mouse Brain in Stereotaxic Coordinates and the hippocampus was cut into 1mm^3^ blocks(2 blocks,respectively). These slabs can store in 25% glutaraldehyde at 4°C for 3 month. Slabs were washed in 0.1 M PBS three times for 15 min, then treated with 1% OsO_4_ for 2h at 4°C dehydrated in escalating concentrations of acetone and flat embedded in Araldite. Ultrathin sections were cut on a microtome, placed onto copper grids, stained with uranyl acetate and lead citrate and examined on a H-7700 microscope. Fifteen distinct regions of the hippocampus were imaged per animal. Images were used to analysis the number/length/width of synapses or the width of synaptic cleft blind to the genotype. A synapse was defined as an electron-dense post-synaptic density area juxtaposed to a presynaptic terminal filled with synaptic vesicles.

### Analysis of apoptosis by FACS

Apoptosis assays on wild type and sh-WDR45 cell lines were performed using Annexin V-FITC apoptosis detection Kit (Beyotime, #C1062, CHN) according to the manufacturer’s protocol. Numbers of annexin V+/PI- and annexin V+/PI+ cells were detected using BD FACSCalibur and data were processed with FlowJo program.

### TUNEL assay for apoptosis

Frozen brain sections from WT and KO mice (16 months) were stained with TUNEL kit (Yeasen, #403006, CHN) according to manufacturer’s instructions. Pictures were acquired using an Olympus-KCC-REM microscope.

### Quantitative proteomics

Mouse brains 6-8 months of age were dissected into PFC (prefrontal cortex), HIP (hippocampus) and MIB (middle brain). Each sample was lysed in urea buffer (7 M urea, 2 M thiourea, 50 mM Tris-HCl, 150 mM NaCl, 1 mM EDTA, pH 7.5, with protease and phosphatase inhibitors). For TMT labeling, 35 ug protein lysate per sample were reduced with 10mM DTT at 55°C for 30 min, alkylated with 50 mM iodoacetamide in the dark for 30 min, precipitated with six volumes of pre-chilled acetone overnight, and digested with trypsin (1:100, Promega) overnight at 37°C in 100 mM TEAB. Peptides from the ten conditions were labeled with 0.8 mg of 10xPlex TMT (Thermo Scientific, #90110) respectively, and combined afterward following the manufacturer’s instructions.

A total of 350 μg tryptic peptides in each brain region was separated into 60 fractions using high-pH RPLC (Waters XBridge C18 column 5 μm, 250 mm × 4.6 mm; mobile phase A (5 mM NH_4_COOH, 2% acetonitrile, pH = 10.0) and B (5 mM NH4COOH, 98% acetonitrile, pH = 10.0)) at a flow rate of 1 ml/min using the following linear gradient: 0-6% phase B for 6 min, 6%-28.5% phase B for 30 min, 28.5%-34% phase B for 5 min, 34%-60% phase B for 10 min, 60% phase B for 9 min. The eluate was collected 1 minute into vials (1 ml/tube). Finally, 60 fractions were combined into 12 components in zigzag fashion. Peptides were concentrated in a speed Vac, desalted in C18 STAGE-tips and primed for LC-MS analysis.

Peptides were analyzed on an EASY-nLC1000 LC (Thermo Scientific) coupled on line with the Q-Exactive Orbitrap mass spectrometer (Thermo Scientific). Peptides were separated on an in-house packed C18-column (15 cm, 75 μm I.D., 3 μm p.s.) with a 120-minutes gradient from 12% to 32% buffer B (98% ACN, 0.1% formic acid) at a flow rate of 250 nl/min. One full scan MS from 400 to 1600 m/z followed by 12 MS/MS scan were continuously acquired. The resolution for MS was set to 70000 and for data-dependent MS/MS was set to 35000. The nano-electrospray source conditions were: spray voltage, 1.8 KV; no sheath and auxiliary gas flow; heated capillary temperature, 275°C. For HCD, the isolation window was set to 2 m/z and the normalized collision energy of 32% was applied.

### Analysis of mass spectrometry data

The MaxQuant (version 1.4.1.2) software was used to analyze the MS/MS raw data. For the database searching parameters, the precursor mass tolerance was set to 15 ppm. Trypsin/P was set as the protease, accounting for in-source fragmentation of lysine or arginine residues followed by proline. Two missed cleavages were allowed. All data were searching against with the UniPort Mouse database (sequences) including oxidation (+15.9949 Da) of methionine, and TMT tenplex (+229.1629 Da) on lysine and carbamidomethylation (+57.0215 Da) of cysteine as static modifications.

### Parallel Reaction Monitoring

Protein candidates with more than 2-fold change and adjust *P*-value of less than 0.05 were selected for further validation by targeted liquid chromatography-parallel reaction monitoring (LC-PRM) MS. Peptides were separated on C18-column with a 75-minute gradient from 12% to 32% buffer B (98% ACN, 0.1% formic acid) at a flow rate of 250 nl/min. PRM MS2 spectra were collected at a resolution of 17500 with AGC target value of 2e^4^. Raw data were analyzed by Skyline using a data-independent acquisition method (version 3.6.1), with a q value of 0.1 (MacLean, Tomazela et al., 2010).

### Bioinformatics analysis

Principal component analysis (PCA) was conducted to demonstrate overall differences of samples with the function princomp in R. Functional annotation was performed with the David bioinformatics resource v6.7 (https://david.ncifcrf.gov/). The functional annotation term categories of biological processes, cellular components and molecular functions are analyzed.

### Statistics

Data are presented as standard error of the mean (SEM) of multiple replicates.All data presented are representative of at least three independent experiments. P-values were generated by two-tailed t-test assuming equal variance using Prism Graphpad Software (San Diego, CA). The value 0.05 (*), 0.01 (**) and 0.001 (***) was assumed as the level of significance for the statistic tests.

### Study approval

All experiments involving animal treatment and care were performed following the Institutional Animal Care and Use Committee protocols from the East China Normal University (ECNU).

*Note: Supplemental Information is available in the online version of this paper. Raw mass spectrometry data is available in PXD008889*

## Reference

Araújo R, Garabal A, Baptista M, Carvalho S, Pinho C, de Sá J, Vasconcelos M (2017) Novel WDR45 mutation causing beta-propeller protein associated neurodegeneration (BPAN) in two monozygotic twins. Journal of Neurology 264: 1020–1022

Bakula D, Müller AJ, Zuleger T, Takacs Z, Franz-Wachtel M, Thost A-K, Brigger D, Tschan MP, Frickey T, Robenek H, Macek B, Proikas-Cezanne T (2017) WIPI3 and WIPI4 β-propellers are scaffolds for LKB1-AMPK-TSC signalling circuits in the control of autophagy. Nature Communications 8: 15637

Behrends C, Sowa ME, Gygi SP, Harper JW (2010) Network organization of the human autophagy system. Nature 466: 68–76

Castro-Caldas M, Carvalho AN, Rodrigues E, Henderson CJ, Wolf CR, Rodrigues CMP, Gama MJ (2012) Tauroursodeoxycholic Acid Prevents MPTP-Induced Dopaminergic Cell Death in a Mouse Model of Parkinson’s Disease. Molecular Neurobiology 46: 475–486

Choi AMK, Ryter SW, Levine B (2013) Autophagy in Human Health and Disease. New England Journal of Medicine 368: 1845–1846

Dall’Armi C, Devereaux KA, Di Paolo G (2013) The Role of Lipids in The Control of Autophagy. Current biology : CB 23: R33–R45

Frakes AE, Dillin A (2017) The UPRER: Sensor and Coordinator of Organismal Homeostasis. Molecular Cell 66: 761–771

Fumagalli F, Noack J, Bergmann TJ, Presmanes EC, Pisoni GB, Fasana E, Fregno I, Galli C, Loi M, Solda T, D’Antuono R, Raimondi A, Jung M, Melnyk A, Schorr S, Schreiber A, Simonelli L, Varani L, Wilson-Zbinden C, Zerbe O et al. (2016) Translocon component Sec62 acts in endoplasmic reticulum turnover during stress recovery. In pp 1173-+-1173-+. Nature Publishing Group

Gregory A, Hayflick SJ (2011) Genetics of Neurodegeneration with Brain Iron Accumulation. Current Neurology and Neuroscience Reports 11: 254–261

Grumati P, Morozzi G, Hölper S, Mari M, Harwardt M-LIE, Yan R, Müller S, Reggiori F, Heilemann M, Dikic I (2017) Full length RTN3 regulates turnover of tubular endoplasmic reticulum via selective autophagy. eLife 6: e25555

Haack Tobias B, Hogarth P, Kruer Michael C, Gregory A, Wieland T, Schwarzmayr T, Graf E, Sanford L, Meyer E, Kara E, Cuno Stephan M, Harik Sami I, Dandu Vasuki H, Nardocci N, Zorzi G, Dunaway T, Tarnopolsky M, Skinner S, Frucht S, Hanspal E et al. (2012) Exome Sequencing Reveals De Novo *WDR45* Mutations Causing a Phenotypically Distinct, X-Linked Dominant Form of NBIA. The American Journal of Human Genetics 91: 1144–1149

Hara T, Nakamura K, Matsui M, Yamamoto A, Nakahara Y, Suzuki-Migishima R, Yokoyama M, Mishima K, Saito I, Okano H, Mizushima N (2006) Suppression of basal autophagy in neural cells causes neurodegenerative disease in mice. Nature 441: 885

Hetz C, Saxena S (2017) ER stress and the unfolded protein response in neurodegeneration. Nature Reviews Neurology 13: 477

Hosp F, Vossfeldt H, Heinig M, Vasiljevic D, Arumughan A, Wyler E, Landthaler M, Hubner N, Wanker Erich E, Lannfelt L, Ingelsson M, Lalowski M, Voigt A, Selbach M (2015) Quantitative Interaction Proteomics of Neurodegenerative Disease Proteins. Cell Reports 11: 1134–1146

Huang DW, Sherman BT, Lempicki RA (2008) Systematic and integrative analysis of large gene lists using DAVID bioinformatics resources. Nature Protocols 4: 44

Huang DW, Sherman BT, Lempicki RA (2009) Bioinformatics enrichment tools: paths toward the comprehensive functional analysis of large gene lists. Nucleic Acids Research 37: 1–13

Kasai-Yoshida E, Kumada S, Yagishita A, Shimoda K, Sato-Shirai I, Hachiya Y, Kurihara E (2013) First video report of static encephalopathy of childhood with neurodegeneration in adulthood. Movement Disorders 28: 397–399

Keene CD, Rodrigues CMP, Eich T, Chhabra MS, Steer CJ, Low WC (2002) Tauroursodeoxycholic acid, a bile acid, is neuroprotective in a transgenic animal model of Huntington’s disease. Proceedings of the National Academy of Sciences 99: 10671

Keestra-Gounder AM, Byndloss MX, Seyffert N, Young BM, Chávez-Arroyo A, Tsai AY, Cevallos SA, Winter MG, Pham OH, Tiffany CR, de Jong MF, Kerrinnes T, Ravindran R, Luciw PA, McSorley SJ, Ba umler AJ, Tsolis RM (2016) NOD1/NOD2 signaling links ER stress with inflammation. Nature 532: 394–397

Khaminets A, Heinrich T, Mari M, Grumati P, Huebner AK, Akutsu M, Liebmann L, Stolz A, Nietzsche S, Koch N, Mauthe M, Katona I, Qualmann B, Weis J, Reggiori F, Kurth I, Hübner CA, Dikic I (2015) Regulation of endoplasmic reticulum turnover by selective autophagy. Nature 522: 354

Kim YC, Guan K-L (2015) mTOR: a pharmacologic target for autophagy regulation. The Journal of Clinical Investigation 125: 25–32

Komatsu M, Waguri S, Chiba T, Murata S, Iwata J-i, Tanida I, Ueno T, Koike M, Uchiyama Y, Kominami E, Tanaka K (2006) Loss of autophagy in the central nervous system causes neurodegeneration in mice. Nature 441: 880

Lu Q, Yang P, Huang X, Hu W, Guo B, Wu F, Lin L, Kovács Attila L, Yu L, Zhang H (2011) The WD40 Repeat PtdIns(3)P-Binding Protein EPG-6 Regulates Progression of Omegasomes to Autophagosomes. Developmental Cell 21: 343–357

MacLean B, Tomazela DM, Shulman N, Chambers M, Finney GL, Frewen B, Kern R, Tabb DL, Liebler DC, MacCoss MJ (2010) Skyline: an open source document editor for creating and analyzing targeted proteomics experiments. Bioinformatics 26: 966–968

Menzies FM, Fleming A, Caricasole A, Bento CF, Andrews SP, Ashkenazi A, Fullgrabe J, Jackson A, Jimenez Sanchez M, Karabiyik C, Licitra F, Lopez Ramirez A, Pavel M, Puri C, Renna M, Ricketts T, Schlotawa L, Vicinanza M, Won H, Zhu Y et al. (2017) Autophagy and Neurodegeneration: Pathogenic Mechanisms and Therapeutic Opportunities. Neuron 93: 1015–1034

Nunes AF, Amaral JD, Lo AC, Fonseca MB, Viana RJS, Callaerts-Vegh Z, D’Hooge R, Rodrigues CMP (2012) TUDCA, a Bile Acid, Attenuates Amyloid Precursor Protein Processing and Amyloid-β Deposition in APP/PS1 Mice. Molecular Neurobiology 45: 440–454

Menzies FM, Fleming A, Rubinsztein DC (2015) Compromised autophagy and neurodegenerative diseases. Nature Reviews Neuroscience 16: 345

Özcan U, Yilmaz E, Özcan L, Furuhashi M, Vaillancourt E, Smith RO, Görgün CZ, Hotamisligil GS (2006) Chemical Chaperones Reduce ER Stress and Restore Glucose Homeostasis in a Mouse Model of Type 2 Diabetes. Science 313: 1137

Popovic D, Akutsu M, Novak I, Harper JW, Behrends C, Dikic I (2012) Rab GTPase-Activating Proteins in Autophagy: Regulation of Endocytic and Autophagy Pathways by Direct Binding to Human ATG8 Modifiers. Molecular and Cellular Biology 32: 1733–1744

Saitsu H, Nishimura T, Muramatsu K, Kodera H, Kumada S, Sugai K, Kasai-Yoshida E, Sawaura N, Nishida H, Hoshino A, Ryujin F, Yoshioka S, Nishiyama K, Kondo Y, Tsurusaki Y, Nakashima M, Miyake N, Arakawa H, Kato M, Mizushima N et al. (2013) De novo mutations in the autophagy gene WDR45 cause static encephalopathy of childhood with neurodegeneration in adulthood. Nature Genetics 45: 445

Shao Y, Guan Y, Wang L, Qiu Z, Liu M, Chen Y, Wu L, Li Y, Ma X, Liu M, Li D (2014) CRISPR/Cas-mediated genome editing in the rat via direct injection of one-cell embryos. Nature Protocols 9: 2493

Tooze SA, Yoshimori T (2010) The origin of the autophagosomal membrane. Nature Cell Biology 12: 831

Wynn DP, Pulst SM (2017) A novel WDR45 mutation in a patient with β-propeller protein-associated neurodegeneration. Neurology: Genetics 3: e124

Yamamoto A, Yue Z (2014) Autophagy and Its Normal and Pathogenic States in the Brain. Annual Review of Neuroscience 37: 55–78

Zhao YG, Sun L, Miao G, Ji C, Zhao H, Sun H, Miao L, Yoshii SR, Mizushima N, Wang X, Zhang H (2015) The autophagy gene Wdr45/Wipi4 regulates learning and memory function and axonal homeostasis. Autophagy 11: 881–890

Zhou Z, Liu A, Xia S, Leung C, Qi J, Meng Y, Xie W, Park P, Collingridge GL, Jia Z (2018) The C-terminal tails of endogenous GluA1 and GluA2 differentially contribute to hippocampal synaptic plasticity and learning. Nature Neuroscience 21: 50–62

